# Microstructural architecture of the argonauts’ shell-like eggcase reveals the formation logic of a convergent extended phenotype

**DOI:** 10.1101/2025.05.27.656349

**Authors:** Kazuki Hirota, Takenori Sasaki, Taro Yoshimura, Shunsuke Onodera, Hirosuke Hirano, Takeshi Toyama, Masa-aki Yoshida, Davin H.E. Setiamarga

## Abstract

Argonautid octopods of the genus *Argonauta* possess a shell-like biomineralized external structure called an eggcase. A classical behavioral observation suggested that this structure is produced by the first dorsal arms rather than by the mantle tissue as seen in typical molluscan shells. In this study, we performed detailed microstructural analyses using scanning electron microscopy to investigate the characteristics of normal, undamaged eggcases and regions that had undergone post-damage repair. Our analysis revealed that the eggcase comprises five layers: an outermost organic membrane, an outer spherulitic-fibrous prismatic layer, a middle organic layer, an inner spherulitic-fibrous prismatic layer, and an innermost organic membrane. Both prismatic layers exhibit bidirectional growth from the middle organic layer, a unique feature not observed in typical molluscan shells. We propose a four-stage formation sequence to explain the observed microstructural architecture: nucleation on the organic scaffold, bidirectional crystal growth from the organic mid-layer, perpendicular growth relative to the eggcase surface, and final encapsulation by an organic membrane. Interestingly, the growth pattern and crystal shapes in the prismatic layers resemble of the internalized shells of cuttlefish, the calcified skeleton of stony corals, and avian eggshells, suggesting functional convergence. We also identified two repair mechanisms: reattachment of broken fragments and regeneration via new secretions. These findings call into question the previous assumptions about the role of the first dorsal arms in calcification. The eggcase also exemplifies the formation of a functionally and structurally complex extended phenotype through convergence.

## Introduction

The argonauts, commonly known as the ‘paper nautilus’, are pelagic octopods belonging to Argonautidae, a family comprising four species: *Argonauta argo*, *A. hians*, *A. nodosus*, and *A. nouryi*^1^, with a secondarily obtained shell-like biomineralized eggcase as an autapomorphy. Together with Alloposidae (e.g. the blob octopus), Ocythoidae (e.g. the football octopus), and Tremoctopodidae (e.g. the blanket octopus), Argonautidae forms Argonautoidea, a superfamily with pelagicity as one of their synapomorphies. Although long thought to be a sister group to the rest of the octopods in Octopodiformes, recent phylogenetic analyses have firmly nested Argonautoidea within Octopodidae^2^. However, unlike their benthic octopod relatives, members of Argonautoidea exhibit traits adapted for life in the open ocean, such as specialized buoyancy control and reproductive strategies.

One key morphological innovation associated with the pelagic lifestyle of the argonauts is the eggcase, a spirally coiled, calcified eggcase that closely resembles the external shells of other mollusks, resembling nautiloid and gastropod shells (Fig. 1). The structure occurs exclusively in females, representing a case of acute sexual dimorphism^1,3^. Similar to the nautilus (*Nautilus pompilius*), an early-diverging extant cephalopod with calcified external shell^4,5^, the shell-like eggcase facilitates neutral buoyancy, thus enabling the holopelagic lifestyle^6^. Argonaut eggcase also serves as a brooding chamber where eggs are laid and attached to its inner surface^7^. In one of the earliest experimental studies on argonaut eggcase formation, Villepreux-Power (1856) used a custom-designed tank to investigate whether the eggcase was a reused structure from another organism or autonomously produced. In her first experiment, she put newly hatched individuals in isolated cages and confirmed that they produced the eggcase spontaneously without any external assistance. In the second, she observed how the argonauts used their first dorsal arms to retrieve and reassemble broken fragments. Based on these observations, she concluded that the eggcase is formed and repaired by the first dorsal arms, unlike the typical, mantle-derived molluscan shell. Additionally, phylogenetic analyses have shown that argonauts diverged from shell-less octopuses (including the rest of the families in Argonautoidea)^2,9,10,11,12^. These results indicate that the eggcase is a product of convergent evolution, rather than a homolog of the typical molluscan shell^13,14,15^.

**Fig. 1.**
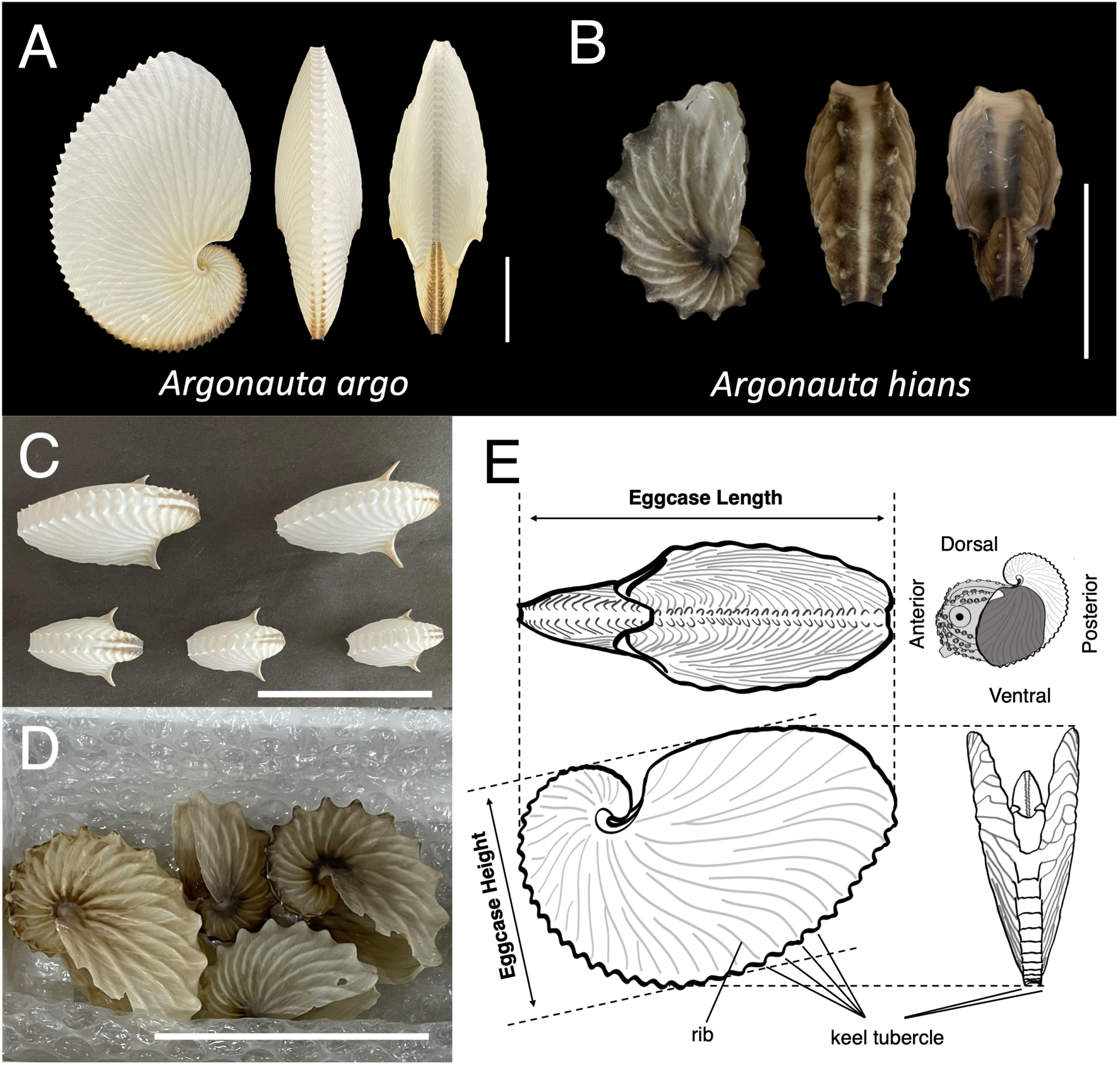
Shell-like eggcases of two argonaut octopuses. A, B) Shell-like eggcases of A) *Argonauta argo* and B) *Argonauta hians*, shown from lateral, posterior, and anterior perspectives. C) Tiny eggcases of *A. argo*. D) Several eggcases of *A. hians* in a container. A range of eggcase sizes was used in this study. E) Schematic diagrams of the living argonaut and its eggcase. All scale bars represent 5 cm.

Microstructural observation of biomineralized structures provides key insights onto their formation mechanisms and evolution^16,17^. Biomineralized structures consist of structurally complex microstructures arranged in layers^18,19^. Each layer exhibits a three-dimensional architecture with specific patterns, formed through a combination of mineral components and organic matrices (e.g., prisms, nacre, foliated, crossed-lamellar). Microstructural observations on several argonaut species^13,14,20,21,22^ have shown that the eggcase is also composed of complex, layered architectures in which each stratum exhibits a specific three-dimensional organization formed by the interactions between mineral elements and organic matrices. These studies also found that microstructurally, the argonaut eggcase generally consists of three layers: an outer spherulitic-fibrous prismatic layer, a middle organic layer, and an inner spherulitic-fibrous prismatic layer. These calcified layers grow radially from the middle organic layer^21^. Chemically, the argonaut eggcase is primarily composed of high-Mg calcite (∼5.1 wt%) and a high proportion of organic matter (∼1.8%)^14,22^. As one of the primary parts of the organic matter, the organic framework, which envelopes each prismatic crystal, plays a pivotal role in controlling both crystal growth and crystal architecture of the eggcase, presenting the intricate interplay between mineral and organic components in biomineralization^14^.

Microstructural observations revealed that the bidirectional crystal growth from the middle organic layer in the argonaut eggcase is distinct from the unidirectional growth observed in typical molluscan shells^21^. Histological studies further underscored these differences, demonstrating variation in developmental modes^23^. In most molluscan species, the mantle tissue responsible for secreting the shell components originates from a shell gland primordium that forms through the evagination of epidermal cells during embryogenesis^23^. In contrast, the shell gland in argonauts disappears entirely during embryogenesis without undergoing evagination. Female argonauts have no eggcase at hatching, and eggcase formation begins *ca.* 12 days after hatching at a size of *ca.* 5-7 mm mantle length^1,24^.

The lack of comparative microstructure studies on argonaut eggcases and other biomineralized structures limits our understanding of convergent biomineralization. In this study, we investigated the detailed microstructural architecture and formation process of the argonaut eggcase in two argonaut species, the common paper nautilus (*Argonauta argo*, Linnaeus 1758) and the brown paper nautilus (*Argonauta hians*, Dillwyn 1817). For this study, we obtained not only fresh eggcase samples of both species, but also fresh *A. argo* specimens with scars indicating prior breakage and repair. By analyzing these samples, our study aims to re-examine previous experimental observations that argonaut species can produce their eggcases autonomously through the involvement of the first dorsal arms^8^ and to explore the function and adaptive significance of the argonaut eggcase based on its microstructural architecture. Our findings provide new insights into the stepwise formation and functional significance of the spherulitic-fibrous prismatic structure in the argonaut eggcase, offering a perspective on the biomineralization processes underlying the logic of formation mechanisms behind the extended phenotypes seen across different molluscan lineages, be it homologous, or analogous.

## Results

### Morphology and mineralogy of the argonaut eggcase

Female argonauts produce a calcareous eggcase resembling loosely coiled gastropod and cephalopod, although it is mechanically light and brittle, unlike the more robust shells of other mollusks (Fig. 1). In *Argonauta argo*, the eggcase is entirely white, while in *A. hians* it is dark brown to black in the posterior and ventral regions. The eggcase size varies among individuals. In the specimens examined, eggcases of *A. argo* ranged from 2.3 by 3.1 cm to 18.3 by 21.6 cm. Those of *A. hians* ranged from 3.7 by 6.4 cm to 4.7 by 7.4 cm (Table S1; Fig. 1). All eggcases are thin and fragile, becoming thicker and harder toward the center of the spiral. Both species possess keel tubercles, which are small protrusions aligned along the ventral keel in two rows.

However, the arrangement and morphology of these tubercles differ between species. In *A. argo*, keel tubercles with acute tips begin in a slightly alternating pattern that gradually shifts into parallel rows. In *A. hians*, keel tubercles have more obtuse tips and maintain a completely alternating pattern throughout. The surface of the eggcase also shows a rib structure. This feature is species-specific. In *A. argo*, one rib aligns with each keel tubercle. In *A. hians*, no such one-to-one correspondence is observed.

Raman spectrometry was used to identify the mineral composition of the eggcase in *A. argo.* Spectral peaks were detected at 288, 717, and 1,086 cm⁻¹ (Fig. S1). These results confirm that the eggcase is primarily composed of calcite. Measurements of magnesium content in the shell fragments showed less than 10 percent molar substitution. These findings are consistent with previous studies on *A. nodosus*^22,25^ and *A. hians*^14^ (Fig. S2).

### Shell-like eggcase microstructure

#### 1. General microstructure and growth pattern

Despite species differences, the eggcase microstructures of *Argonauta argo* and *A. hians* are highly similar, typically consisting of three layers: an outer spherulitic-fibrous prismatic layer, a middle organic layer, and an inner spherulitic-fibrous prismatic layer (Figs. 2A-C, S3). The crystals in both calcified prismatic layers appear as elongated polygonal prisms with fibrous interconnections, creating a surface appearance resembling a honeycomb. In vertical section, individual crystals appear as elongated prisms measuring between 2 µm and 10 µm in length along the long axis. In surface view, they exhibit n-polygonal cross-sections (n = 3 - *ca.* 20), with an average surface area of approximately 1.0 µm². The crystal growth pattern generally suggests that the two prismatic layers grew bidirectionally from the middle organic layer (Fig. 2C). The radial growth on both sides started from a single point in the middle organic layer, referred to as the crystal nucleation point. Once crystal nucleation occurred, the crystals grew while competing for growing space with neighboring crystals, gradually becoming uniform in size and arrangement in a direction perpendicular to the eggcase surface (Fig. 2D). The height of adjacent crystallites appears to be evenly regulated, and each crystallite’s height seems to increase from the middle region towards the edges (Fig. 2C, D). The surface view indicates that the polygonal outlines of individual crystals often exhibited partial cracking (Fig. 2E).

**Fig. 2.**
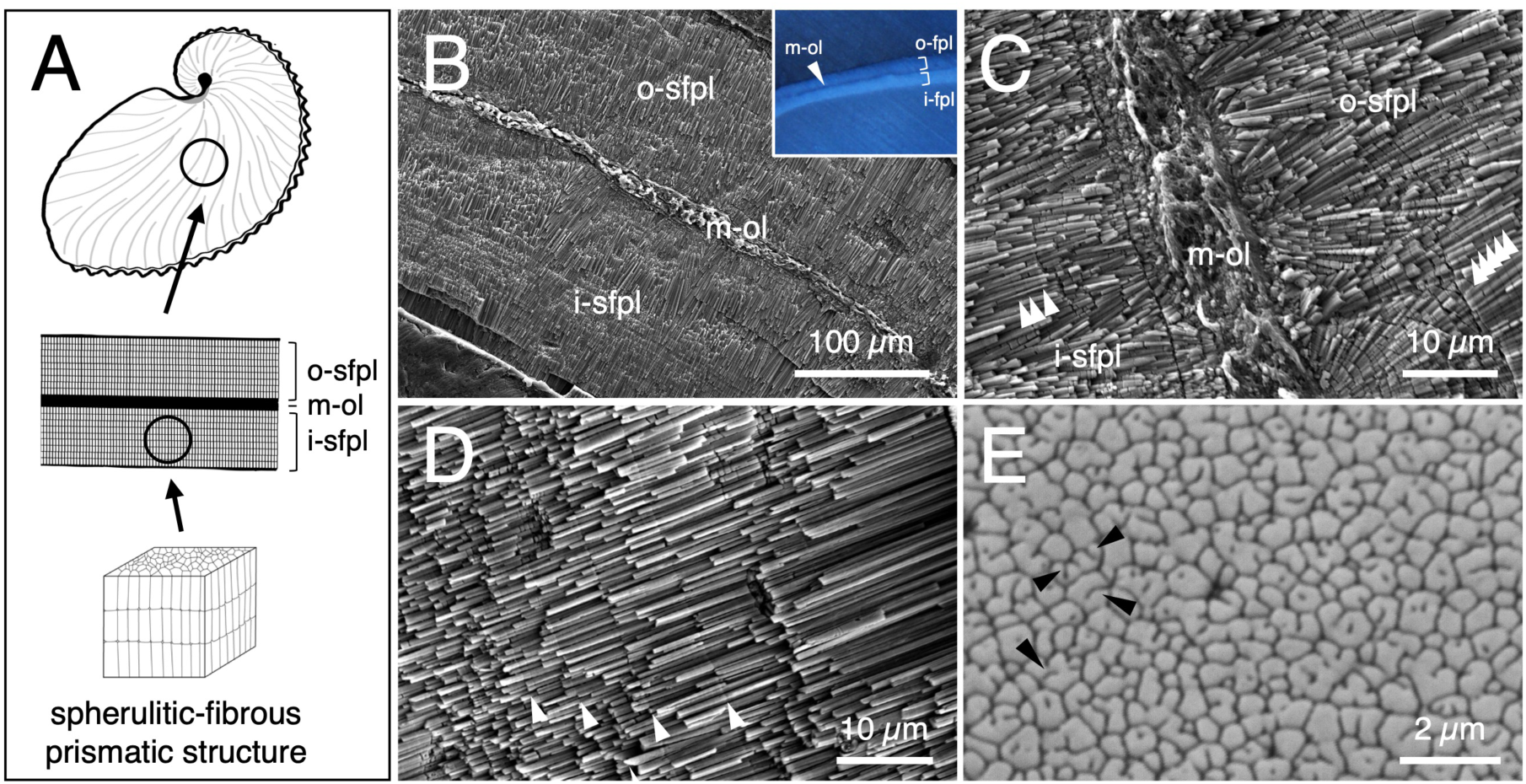
SEM micrographs showing the general microstructure of argonaut eggcase. General microstructure of the *A. argo* eggcase (Fig. S3 for *A. hians*): A) Schematic diagrams of the general microstructure of the eggcase. B) Vertical section of the eggcase. The inset image was taken using stereoscopic microscopy. C, D) Enlarged vertical section from B, representing C) the middle and D) the distal regions, respectively. White arrowheads indicate examples of crystal boundaries. E) Surface view of the prismatic structure in the eggcase. Black arrowheads indicate examples of a partial cracking in the prismatic structure. Abbreviations: outer spherulitic-fibrous prismatic layer (o-sfpl), middle organic layer (m-ol), inner spherulitic-fibrous prismatic layer (i-sfpl)

#### 2. Region-specific microstructural modifications

Vertical section of argonaut eggcase from various locations revealed that localized modifications of the general microstructure give rise to the formation of region-specific features. Two localized regions show modified microstructure: the center of the spiral (Figs. 3A, S3) and the keel tubercles (Figs. 3B, S3). In the center of the spiral, the inner and outer prismatic layers appear connected, forming a closed structure that envelops the middle organic layer (Fig. 3A, C, E, F). In contrast, the keel tubercle region, characterized by its regularly jagged structure along the ventral side, terminates in either an acute or obtuse angle (Fig. 3B, D, G, H). The angular tips form through deformation of the middle organic layer. When the tip is acute, the middle organic layer is split, resulting in a hollow internal gap. When the angle is obtuse, the middle organic layer is simply curved without forming any gap (Fig. S3).

**Fig. 3.**
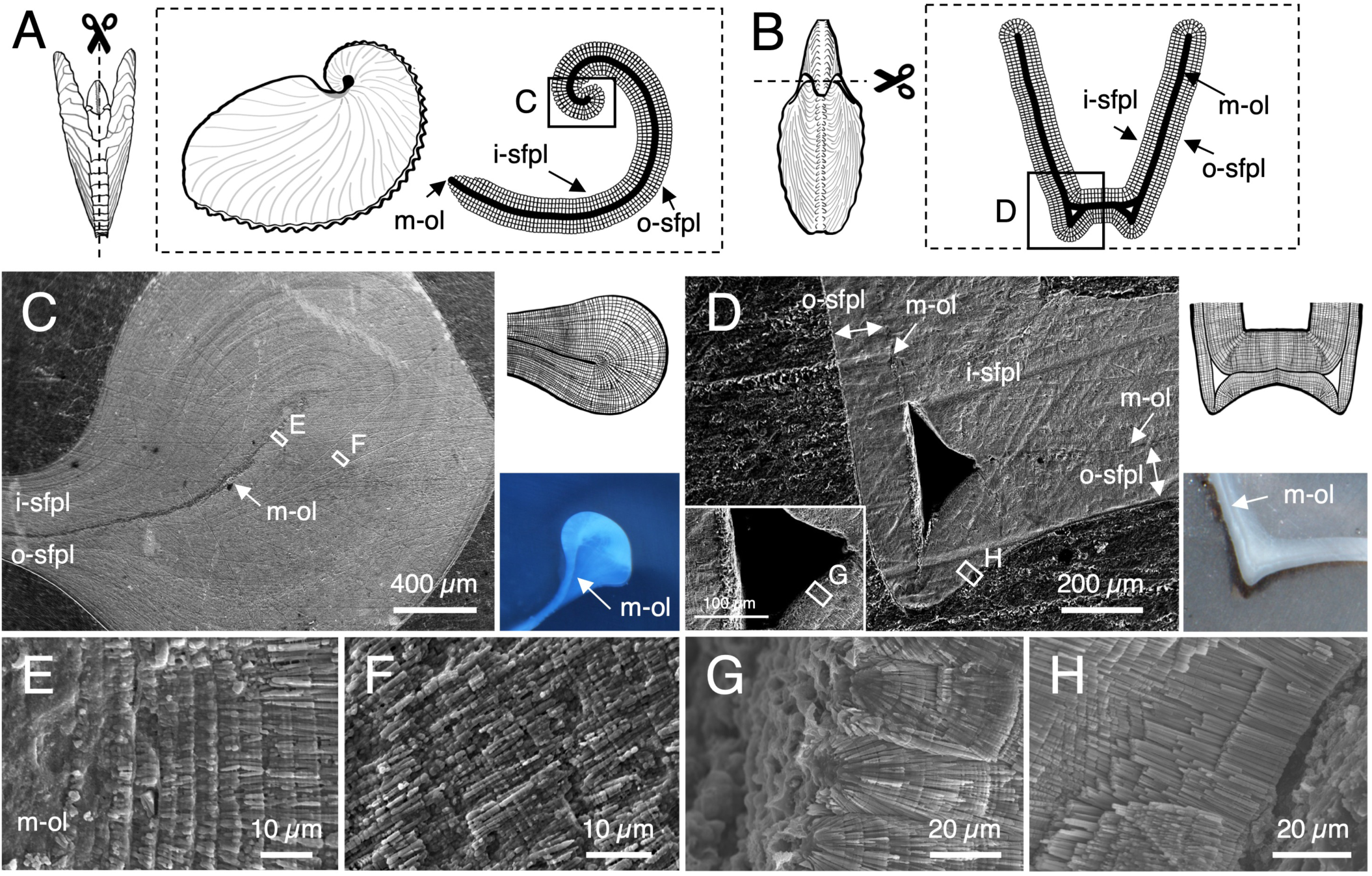
Schematic diagrams, Stereomicroscopic images, and SEM micrographs of vertical sections of *A. argo* eggcase. Schematic diagrams, Stereomicroscopic images, and SEM micrographs showing the *A. argo* eggcase (Fig. S3 for *A. hians*). A, B) Schematic diagrams depicting the overall structure and microstructure of the eggcase, highlighting two local regions: A) the center of the spiral and B) the keel tubercle. C, D) SEM images, schematic diagrams, and optical microscope images of the microstructure at C) the center of the spiral and D) the keel tubercle. E, F) Enlarged views of the center of the spiral from C. G, H) Enlarged views of the keel tubercle from D. Abbreviations: outer spherulitic-fibrous prismatic layer (o-sfpl), middle organic layer (m-ol), inner spherulitic-fibrous prismatic layer (i-sfpl)

#### 3. Region-specific interior and exterior surface structures

We conducted detailed observations on the microstructures across marginal to central regions of both the interior and exterior surfaces of the eggcase. While the general surface microstructure was observed ubiquitously (Fig. 2E), crystal-growing surfaces were distributed non-uniformly (Figs. 4, S4). At the margin of the interior surface, where crystal nucleation occurs, high crystal growth activity was observed. Crystal growth varied mosaically between individuals: some regions were in an active growth phase (Fig. 4A), while others were in a non-growth phase (Fig. 2E). At the periphery of the nucleation sites, crystallites were extremely irregular in size and orientation, and organic matter was more abundant around them. The interior surface displayed concave-convex patterns based on the height of crystallites, which formed clustered assemblies (Fig. 4A). The borders at both ends of the elongated crystallites contained abundant organic sheets, corresponding to the initial crystallization floor (ICF; Fig. 4A1) and the crystallization termination floor (CTF; Fig. 4A1). Toward the middle area of the interior surface, most areas exhibited a decrease in crystal growth activity and in the amount of organic deposition (Fig. 4B). Occasionally, crystal growth was observed in a patchy manner. Unlike the crystal growth pattern observed in the margin area, the middle area showed gel-like aggregates with low organic content on the horizontal surfaces of the crystallites (Fig. 4B1). The interior surface retained concave-convex morphology at the level of the individual crystallites (Fig. 4B2). The crystal growth pattern observed from the vertical section, which shows crystallites transitioning from random to organized perpendicularly to the eggcase surface (Fig. 2), was broadly consistent with the characteristics observed from the marginal to the middle area. In the central region of the interior surface, no ongoing crystal growth was observed. Instead, the crystal surfaces were covered with an organic membrane in a mosaic pattern (Fig. 4C). The shape of the organic membranes resembled those found in the margin area when the crystallites were covered at the termination of crystal growth. The organic membrane was thin and slightly translucent, beneath which the prismatic structure was faintly visible.

**Fig. 4.**
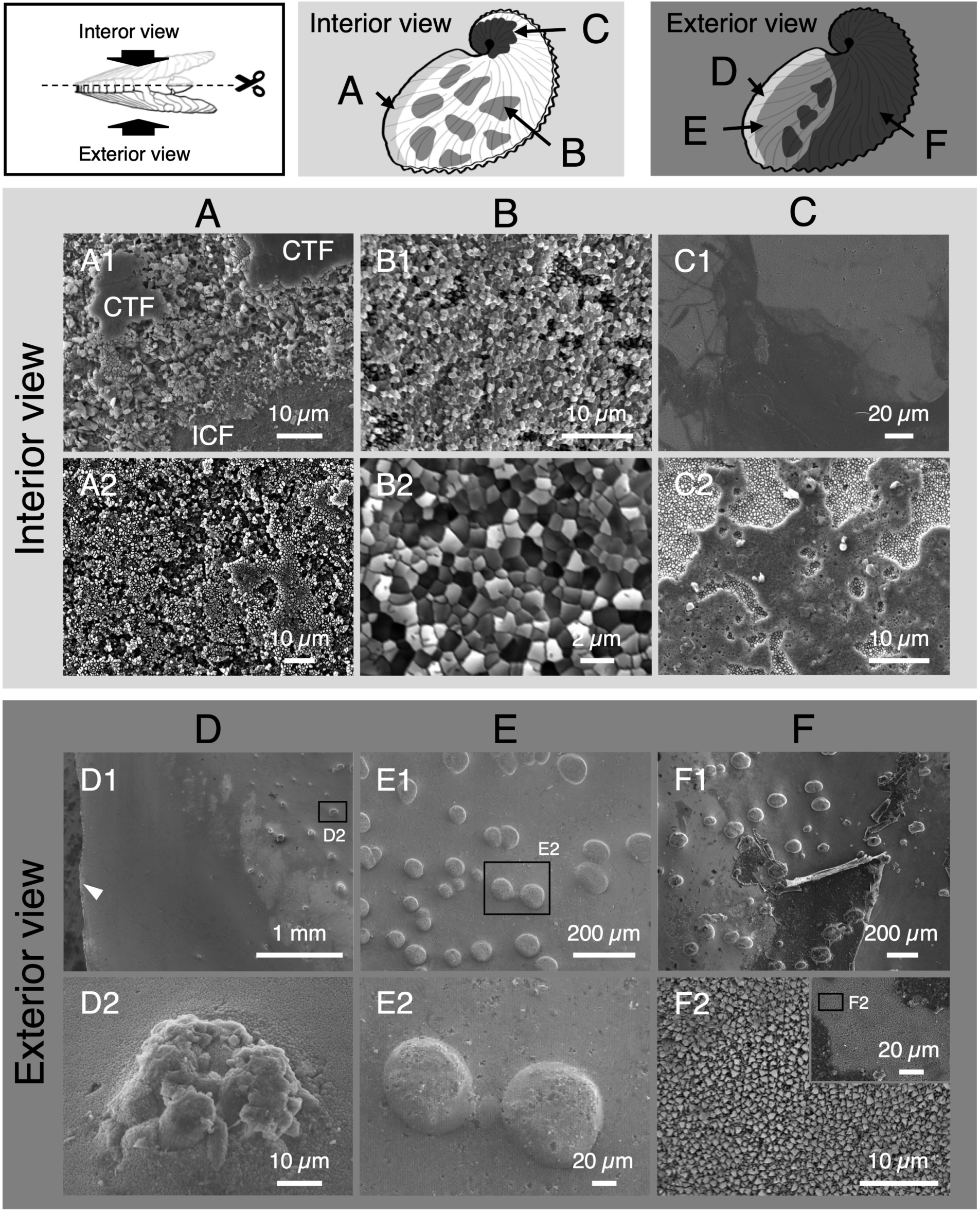
Schematic diagrams and SEM micrographs of surface view of *A. argo* eggcase. Schematic diagrams and SEM micrographs showing the surface view of the microstructure of the *A. argo* eggcase (Fig. S4, S5 for *A. hians*). All samples for surface observation were unetched. The microstructure of A-C) the interior and D-F) exterior surfaces of the eggcase were observed in three regions of A, D) marginal, B, E) middle, and C, F) central regions. White arrowheads in D1 indicate the eggcase aperture. SEM micrographs of vertical sections of the tuberculate micro-ornamentation were shown in Fig. S6. Abbreviations: initial crystallization floor (ICF), crystallization termination floor (CTF)

The exterior surface was characterized by tuberculate micro-ornamentation and a thick organic membrane compared to the characteristics of the interior surface (Figs. 4D-F, S4, S5, S6). The tubercles could be seen from *ca.* 2 mm inward from the margin (Fig. 4D1). These tubercles did not form well-defined hemispheres on the surface; instead, they formed a rough texture with aggregates of crystallites (Fig. 4D2). In the middle region, the exterior surface exhibited well-defined hemispheres (Fig. 4E). The central area including tuberculate ornamentation was completely covered by organic membranes (Fig. 4F). In some areas where the organic layer had rubbed off due to drying, multiple layers of organic material were visible, with crystallites observable beneath them (Fig. 4F2). The surface of each crystallite was sharp. The extent of the organic membrane exhibited a mosaic-like pattern and showed significant individual variation, probably due to secondary factors such as drying. Consequently, the exterior surface exhibits prominent tubercles and thicker organic membranes, characteristics which are distinctly absent from the interior surface.

### Structural repair and regeneration of the eggcase in A. argo

In our specimens, the repair scars were located at the posterior edges (keel) of the eggcase and far from the aperture (Fig. 5). Visual inspection of the microstructures of these scars allowed us to identify two distinct types of regenerated hard tissues (Fig. 5A, B). One type involves fragment adhesion to the interior of the eggcase by an organic layer (“fragment-inserted repair”; FIR). The second type involves filling holes caused by perforation (“hole-sealing repair”; HSR). FIR is characterized by fragments with angular sharpness partially fitted into the broken areas like puzzle pieces (Fig. 5B). These fragments were then covered by an organic membrane to firmly adhere them to the inner surface of the eggcase (Fig. 5B, C). Unsurprisingly, the vertical section of an inserted fragment showed a microstructure consisting of three layers: an outer spherulitic-fibrous prismatic layer, a middle organic layer, and an inner spherulitic-fibrous prismatic layer, which is exactly the same as the layered structure of a normal eggcase (Fig. 5C).

**Fig. 5.**
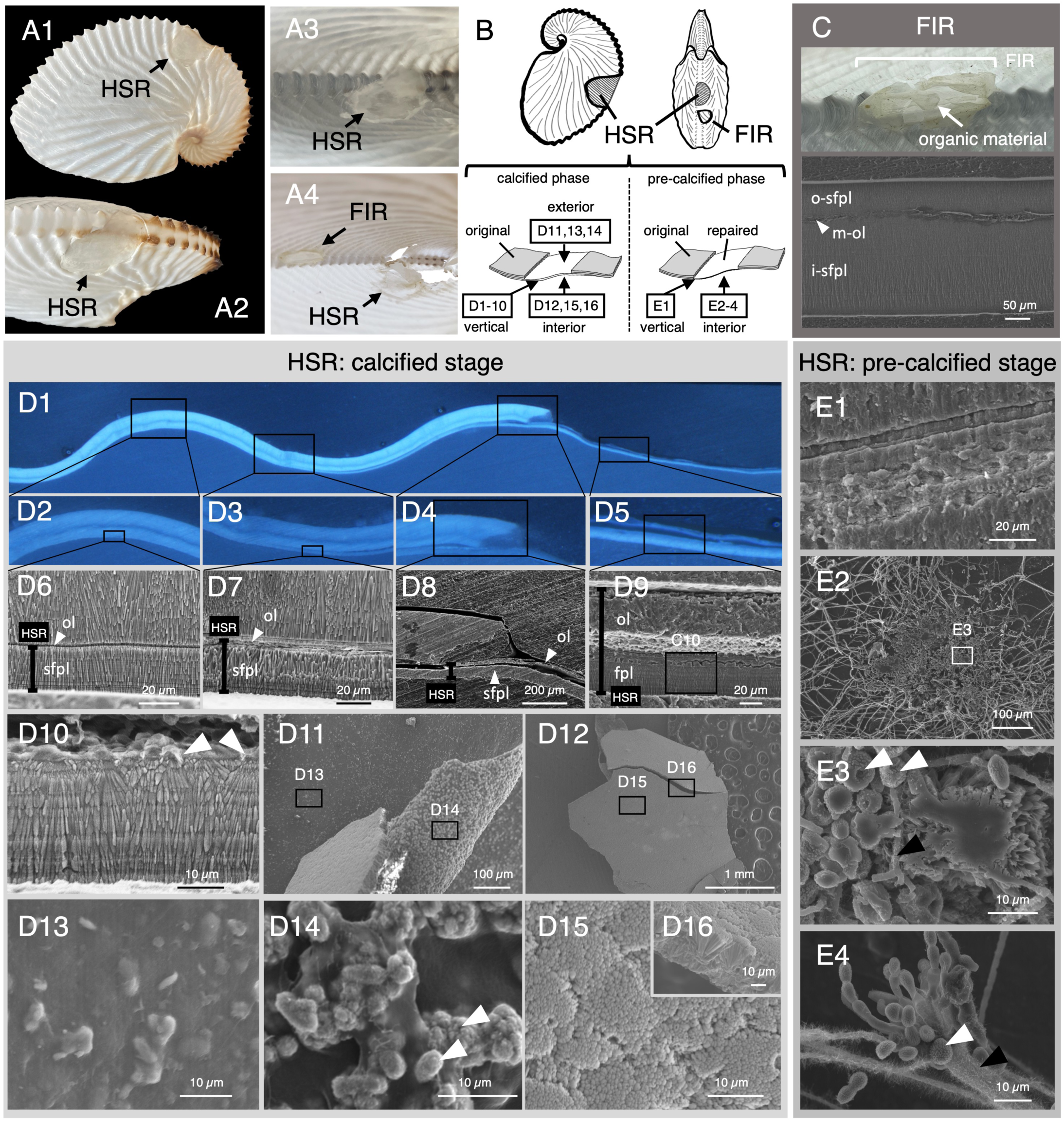
Schematic diagrams, Stereomicroscopic images, and SEM micrographs of repair traits of *A. argo* eggcase. A) Photographs of *A. argo* eggcases with repair traits on the keel region (Table S1). Two distinct biological repair traits were identified: fragments adhering to the interior surface of the eggcase (FIR), and a newly formed structure sealing the broken hole (HSR). B) Illustrations of FIR and HSR. C) Photographs of the HSR, and SEM micrographs of a vertical section of the HSR. D) Stereomicroscopic and SEM micrographs of the HSR in the calcified region. D1-D10) vertical sections; D11, D13, D14) exterior views; D12, D15, D16) interior surface views. D10 is an enlarged view of D9. D14 indicates a region where the organic membrane is rolled or folded. E) SEM micrographs of the HSR in the pre-calcified region. E1) vertical section; E2-E4) interior surface views. White arrowheads of D10, D14, E3 and E4 indicate a spherical particle for the nucleation site. Black arrowheads of E3 and E4 indicate organic fibrils. Abbreviations: outer spherulitic-fibrous prismatic layer (o-sfpl), middle organic layer (m-ol), inner spherulitic-fibrous prismatic layer (i-sfpl), spherulitic-fibrous prismatic layer (sfpl), organic layer (ol), fragment-inserted repair (FIR), hole-sealing repair (HSR)

Through our observation, we identified two distinct phases of HSR: a calcified stage and a pre-calcified stage, each corresponding to a different region. In our samples, the calcified region of the HSR scar did not fully replicate the microstructural layers of a normal eggcase. This region was composed of two layers: an outer organic layer and an inner spherulitic-fibrous prismatic layer, lining and reinforcing the gap/hole from the inside (Fig. 5D1, D5, D9). In areas where the HSR layer was completely separated from the non-repaired regions of the eggcase, crystal growth began with initial crystal nucleation in the spherical particles, followed by radial growth from each particle (Fig. 5D10). The calcified prismatic layer in these regions ranged from 20 µm to 40 µm in thickness. Further observation of the interior and exterior surfaces of the repaired region showed that the outer organic layer was convex, with peeled areas containing spherical particles composed of organic-inorganic complexes (*ca.* 5 µm in diameter; Fig. 5D11-15). These spherical particles, observed in vertical section perpendicular to the surface of the repaired eggcase, correspond to the initial crystal nucleation points for radial growth in the calcified layer (Fig. 5D10). The prismatic layer in the repaired region appears to terminate at the radial growth stage and does not exhibit the subsequent layered growth perpendicular to the eggcase surface. The interior surface of the eggcase displayed shallow hemispherical convexities, each composed of a cluster of crystallites radiating from a single nucleation point (Fig. 5D15, 5D16). The interface between the regenerated and the non-broken regions exhibited local variations in microstructure, reflecting spatial changes from the edge of the gap/hole outward (Fig. 5D1, D4, D8). Meanwhile, in the regions adjacent to the gap/hole, two eggcase layers overlap: that of the original eggcase, and a newly formed HSR layer (Fig. 5D1, D3, D7). The new HSR layer was formed in direct contact with the original inner prismatic layer, with the crystal layer growing radially inward across an organic sheet. In contrast, farther from the gap/hole, the calcified layer retained the granular orientation of the original eggcase, even though it was separated from the layer of the original unbroken eggcase by a thin organic layer (Fig. 5D1, D2, D6). This region is characterized by the absence of crystal nucleation points and a thinner organic layer (*ca.* 2 µm) when compared to that in the regions adjacent to the gap/hole (*ca.* 5 µm) (Fig. 5D6, D7). These proximodistal differences in regional growth patterns likely produce mechanical strength at the adhesion site, thus enhancing the repaired structure’s resistance to subsequent damage.

**Fig. 6.**
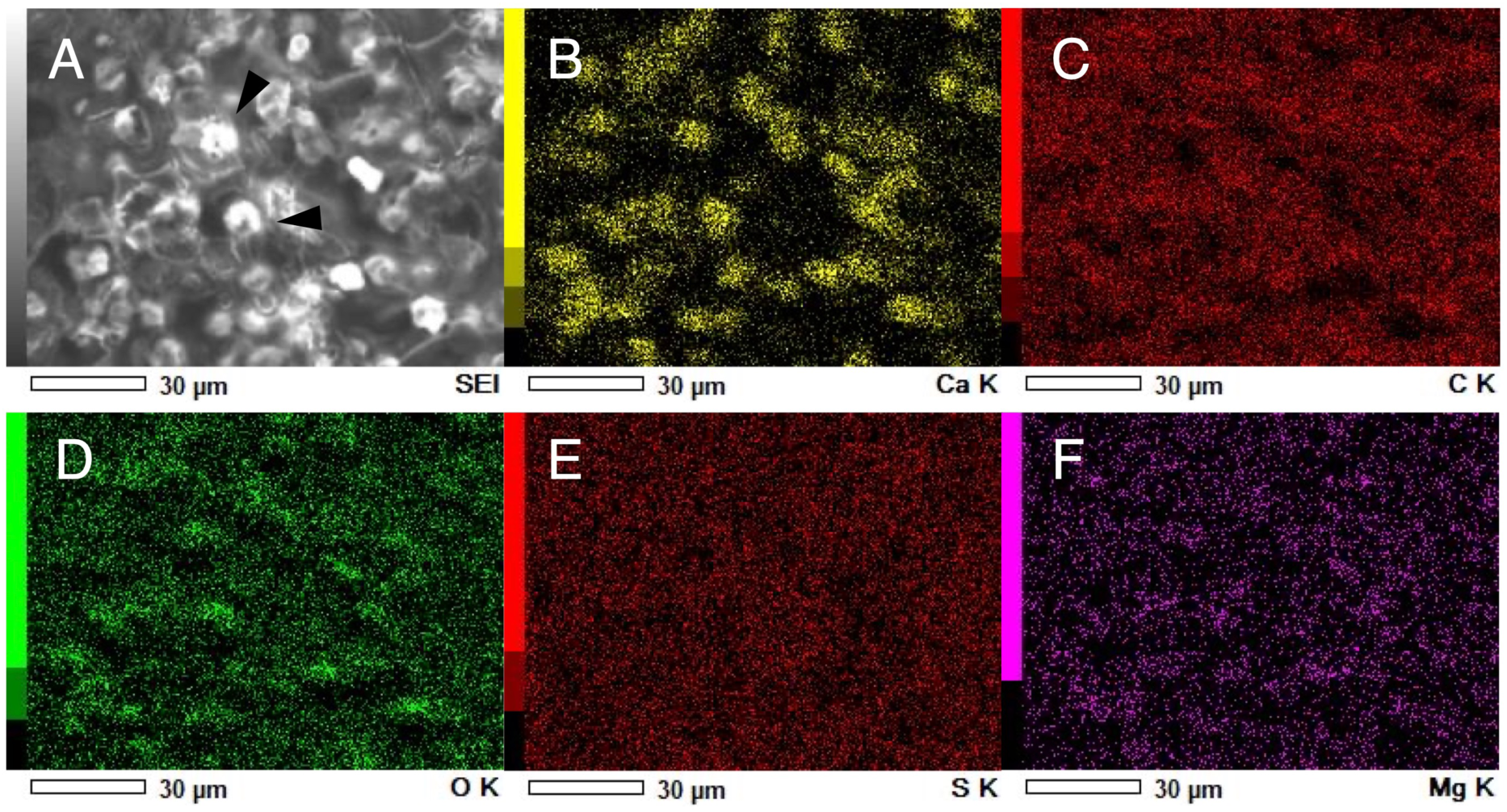
EDS compositional analysis of the original and repaired eggcase of *A. argo*. EDS elemental maps of the repaired *A. argo* eggcase, showing a surface view of the HSR in the pre-calcified region (Fig. 5D14, E3, E4). A) Spherical particles observed on the interior surface of the repaired eggcase. White arrowheads indicate a spherical particle. B-F) Elemental maps of B) calcium, C) carbon, D) oxygen, E) sulfur, and F) magnesium. The calcium distribution reflects the presence of the calcified spherical particles. The particles on the HSR consist of crystals that serve as nucleation sites from which the calcified layer subsequently grows (Fig. 5). No notable differences in elemental composition were observed between the original and repaired eggcases (Fig. S2).

**Fig. 7.**
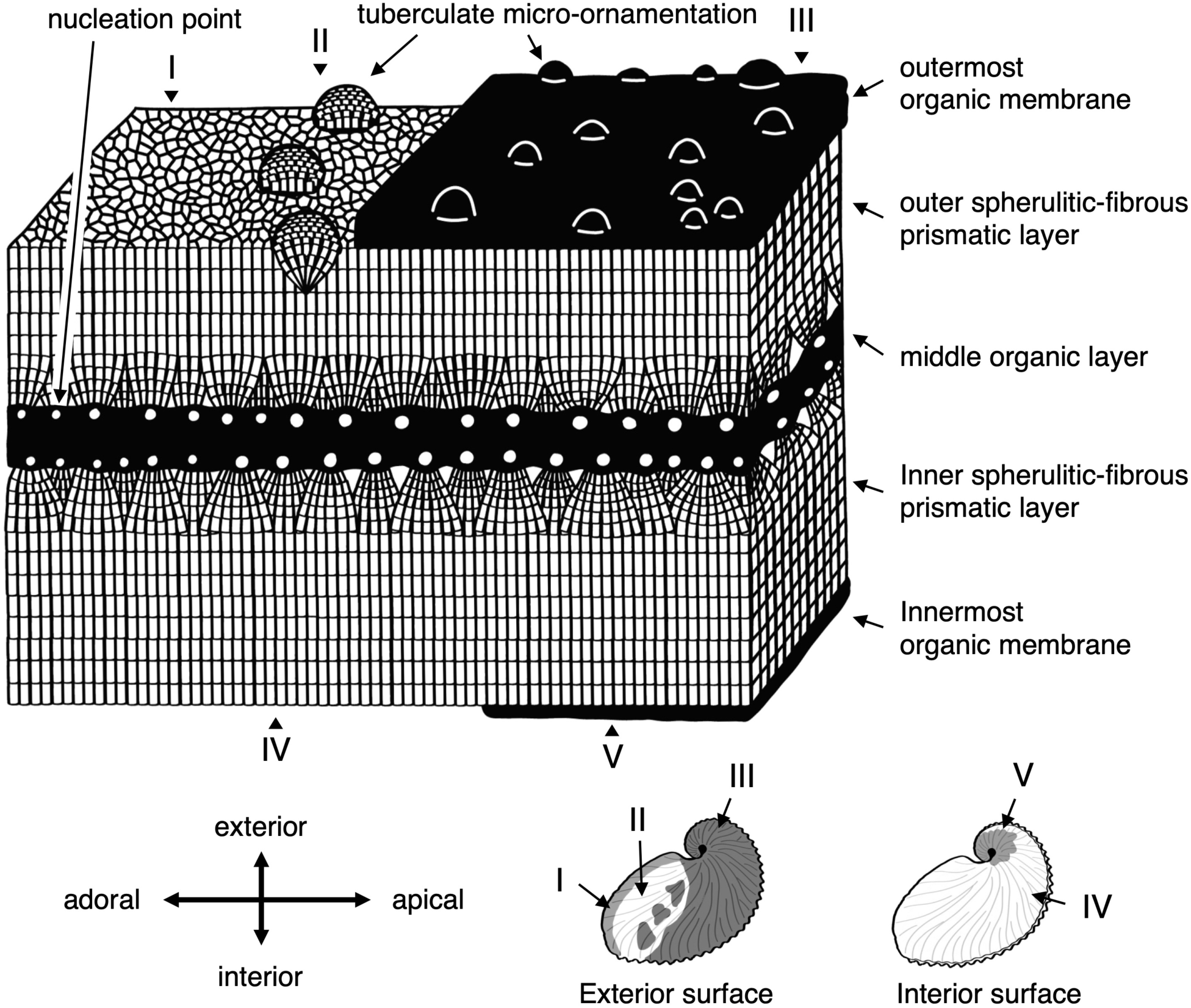
Schematic diagram of the argonauts’ eggcase microstructure. The distribution of the organic membrane and the tuberculate micro-ornamentation is shown at the bottom right. The exterior surface is classified into three types: (I) without the organic membrane and micro-ornamentation, (II) with micro-ornamentation only, and (III) with both the organic membrane and micro-ornamentation. The interior surface is categorized into two types: (IV) without the organic membrane and (V) with the organic membrane.

**Fig. 8.**
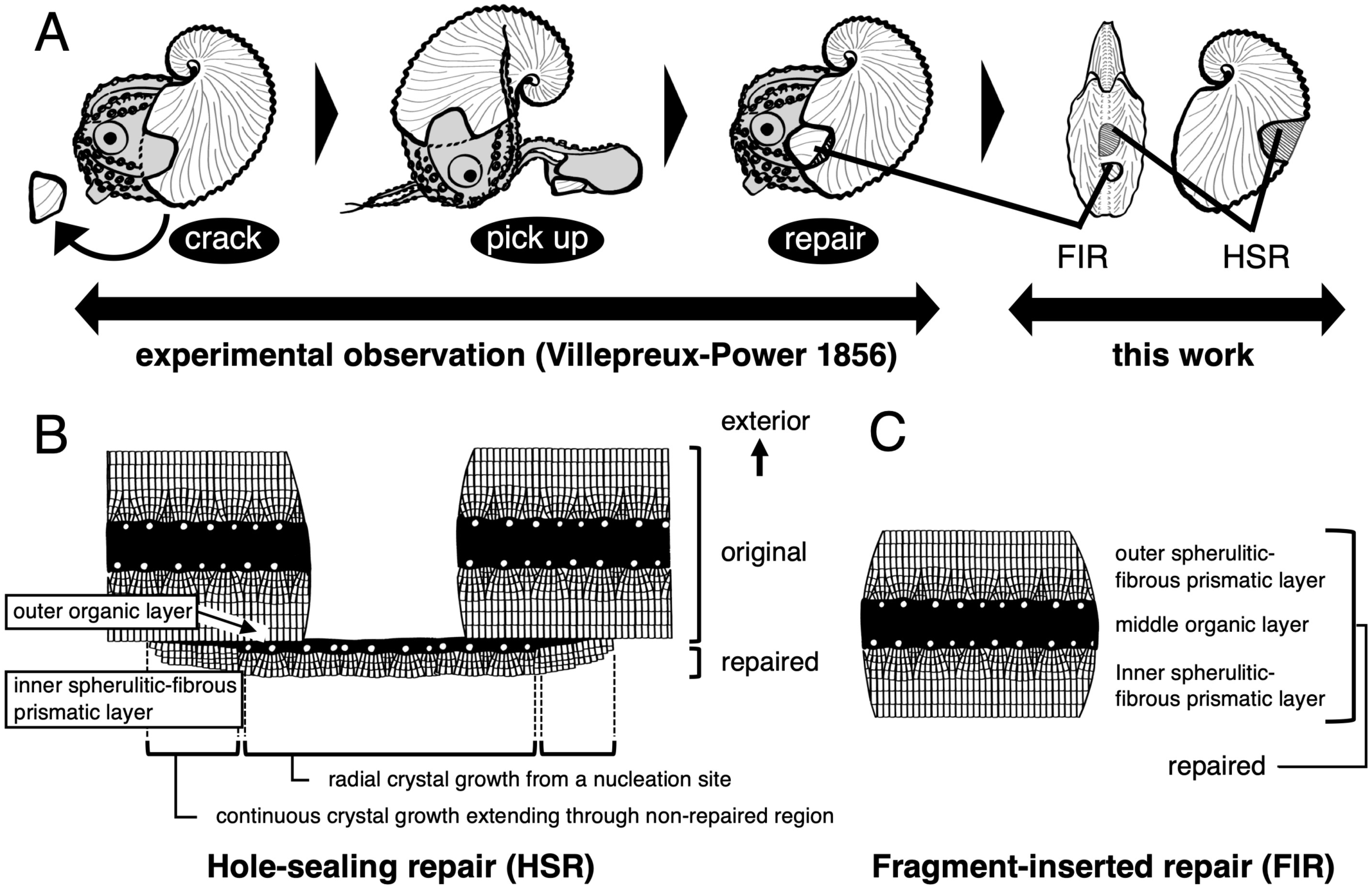
Diagram of the microstructure of repaired argonaut eggcase. A) Schematic illustration of the repair traits of the eggcase. Two types of repair traits are confirmed: hole-sealing repair (HSR) and fragment-inserted repair (FIR). B, C) Diagram of the microstructure of B) the HSR and C) the FIR. Abbreviations: fragment-inserted repair (FIR), hole-sealing repair (HSR)

In contrast to the calcified regions described above, our observations of the pre-calcified stage of HSR revealed the initial phase of crystallization, characterized by spherical particles and assemblages of organic sheet and fibrils (Fig. 5E). The interior surface of this stage exhibited spherical particles and organic fibers. Elemental mapping indicated the presence of calcium in the spherical particles, but not in the surrounding organic sheets (Fig. 6). These spherical particles, likely function as nucleation sites, were deposited around the organic fibers, with some areas displaying spike-like structures radiating from these crystallites (Figs. 5E2,3, S7).

## Discussion

### New findings and functional significance of the argonaut eggcase microstructure

The argonaut eggcase exhibits a three-layered structure, with a middle organic layer sandwiched between outer and inner spherulitic-fibrous prismatic layers^13,14,20,21^ (Fig. 7). Localized modifications occur at the keel tubercles, where the middle organic layer splits into two sheets (Fig. 3A), and at the spiral center, where continuous prismatic layers enclose the middle organic layer (Fig. 3B, 3C, 3E, 3F). The presence of two sheets separated by a gap in the keel tubercles suggests the middle organic layer is a single structural unit composed of two organic sheets, serving as a scaffold for crystal nucleation. Meanwhile, the continuous crystal layer at the center of the spiral suggests that the inner and outer prismatic layers are not only identical in composition and microstructure, but also follow a similar growth pattern in which crystal layers can grow omnidirectionally when an organic layer is present (Fig. 3). These local structural variations indicate that the eggcase adjusts its mechanical strength by altering the shape of the middle organic layer.

Surface observations showed structural contrasts between the inner and outer layers, including differences in organic membrane distribution and tuberculate ornamentation (Fig. 7). Both the innermost and outermost organic membranes were present on the interior and exterior surfaces, respectively, but differed in thickness, texture, and extent. The outermost membrane, composed of stacked organic sheets, was thicker and covered nearly the entire exterior except the adoral margin, while the thinner innermost membrane was confined to the central interior. These structural differences likely reflect important functional adaptations. The inner surface of the inner layer is covered by a thinner innermost organic membrane. We suggest that this innermost organic membrane acted as the termination point for crystal growth, because it resembles the boundary region (the margin) where crystal growth ends, which is visible as organic membranes within the crystals. The localized distribution of the innermost organic membrane and the function of the surrounding region in egg attachment to the surface of the eggcase^26^ suggest that this membrane may serve as an adhesive surface for eggs.

Our observation indicates that the outermost organic membrane covers the entire eggcase and is specifically thick in the posterior areas inaccessible to the first dorsal arm. The areas where the covering organic membrane is damaged and the underlying crystal layers are exposed display lath-like crystallites, which is interpreted as indicative of partial crystal dissolution^22,25^. This suggests that the organic membrane may function to prevent calcium carbonate crystals from dissolving into seawater. In fresh, hydrated samples, the eggcase dissipates mechanical energy through its viscoelastic response that varies with relative humidity and load application rate^27^. In aged, dried, and deteriorated specimens, the outermost organic membrane peels off, causing the eggcase to become brittle. The thick outermost organic membrane likely maintains the eggcase’s flexibility, mechanical strength, and provides protection from dissolution and physical impacts. Known functions of surface-coating organic membranes, such as degradation prevention, adhesion, and mechanical properties enhancement, are also observed in analogous calcareous structures of other organisms, such as coccoliths and eggshells^28,29,30,31,32^.

In addition to the innermost and outermost organic membranes, the eggcase also differs structurally between the inner and outer surfaces. Hemispherical structures 30-150 µm in diameter, known as tuberculate micro-ornamentations, were observed only on the exterior surface of the eggcase and were absent on the interior surface of the eggcase. We suggest that these tubercles may play a role in protection against the impacts from waves and rocks. Therefore, the outer layer of the eggcase forms such a complex architecture to enhance structural strength to prevent damage.

### Comparative microstructural analysis of the argonaut eggcase and other conchiferan shells

The general appearance of the argonaut eggcase superficially resembles that of typical molluscan shells, appearing nearly indistinguishable at first glance. However, microstructural characteristics reveal clear differences in their biomineralization processes. Molluscan shells are secreted by mantle tissue, with mineral deposition proceeding inward from an organic sheet called the periostracum^33^. In contrast, the argonaut eggcase is formed probably by secretions from the first dorsal arm pairs (*sensu* Villepreux-Power), with crystals growing bidirectionally from a middle organic layer. This growth pattern differs markedly from that of typical molluscan shells, in which a calcified layer grows unidirectionally inward from the periostracum^21^.

Among modern cephalopods, the argonaut eggcase may be interpreted as a non-homologous shell-like structure, differing in crystal polymorph rather than microstructure, since modern cephalopod shells also grow bi-directionally from an organic layer^34,35,36^. Unlike primitive cephalopods (e.g., nautilus and ammonite shells), modern cephalopods have modified their shells into internal or non-calcified structures (e.g., cuttlebone, internal shell of *Spirula*). Despite variation in shell macro- and microstructure, all modern cephalopod shells contain aragonite as their carbonate crystal polymorph^37,38,39^. In contrast, the argonaut eggcase consists entirely of calcite, as previously reported^14,21,22^. The selection of crystal polymorphs in biominerals is thought to reflect paleoenvironmental ocean compositions (Mg/Ca ratio) when the biomineral structures were acquired^40^. The origin of cephalopod shells can be traced back to a single origin acquired prior to the Ordovician period, when the Mg/Ca ratio indicated an aragonite sea. Meanwhile, argonauts appeared during the Early Tertiary^10,41^, when the Mg/Ca ratio indicated a calcite sea and ocean chemical compositions predominantly favored calcite^42^. These contrasting crystal polymorphs suggest that the biomineralized structures in cephalopods and argonauts arose independently within the cephalopod lineage.

Comparison of the argonaut eggcase with cephalopod shells at the microstructural level reveals three critical similarities: microstructural architecture, bidirectional growth, and tuberculate micro-ornamentation. The outer calcified layers of the internalized shells in cuttlefish and *Spirula* display a microstructural architecture characterized by the presence of spherulitic prismatic layers^35,36,43^, closely resembling that of the argonaut eggcase. In these species, the spherulitic prismatic layer grows outward from an organic layer (probably periostracum^44^) forming a microstructure with a middle organic layer sandwiched between two calcified layers. This organization is also seen in embryonic/juvenile shells of Mesozoic ammonites^35,36,43,45^. However, despite the resemblance between spherulitic prismatic and layered structures, the microstructure of other cephalopod shells is fundamentally different (i.e., not homologous) from that of the argonaut eggcase. This is because the inner and outer calcified layers in their shells are not continuous and therefore structurally distinct, whereas the argonaut eggcase exhibits a consistent microstructure across both layers. Thus, while the overall architecture and bidirectional growth appear similar, the underlying formation mechanisms differ. These shared microstructural traits may reflect developmental or functional constraints and likely represent a case of convergent evolution within the cephalopod lineage.

Tuberculate micro-ornamentation restricted to the exterior surface has been reported in Mesozoic ammonites and some gastropods^46,47,48,49^. The formation process of tubercles in these groups remains poorly understood. Hickman (2004) proposed that such structures may result from mineralization under weak biological control, based on observations of tubercles in gastropod larval shell. In contrast to these uncertain primordia, our observations indicate that tubercles in the argonaut eggcase develop progressively and are fully developed in the marginal area under tightly regulated biological control. Landman (1994) suggested that the ornamental tubercles in embryonic or juvenile ammonites may aid hatching by facilitating egg capsule rupture. In argonauts, we propose that tubercles help protect the eggcase from mechanical impact caused by waves or contact with rocks, probably similar to gastropods^48^. This interpretation is supported by their widespread distribution on the outer surface of adult eggcase specimens. Although mechanical testing has yet to be performed to confirm their role, the ornamental tubercles may also enhance shell strength before and after hatching.

### Unique mode of eggcase repair in argonauts

Mollusks repair damaged shells by secreting new material from mantle tissue at the injury site. When damage occurs beyond the shell edge away from the mantle, the repair mechanism differs from normal shell formation, as evidenced by distinct microstructural and mineralogical features^51,52,53,54^. In the argonaut eggcase, the surface of the repaired region, which was damaged by perforation in the posterior section of the keel, appears optically opaque and smooth, clearly differing from the non-repaired regions (Fig. 8A). Our microstructure observations showed that the repaired area comprises only two layers, the outer organic and the inner calcified layers, lacking the outer calcified layer present in the original three-layered structure (Fig. 8B). Moreover, the outer organic layer does not extend from the middle organic layer but instead is located beneath the inner calcified layer of the intact area. At the adhesion site of repaired and non-repaired regions, we observed a gradual thinning of the organic layer with increasing distance from the perforation. Crystal orientation became increasingly consistent with that observed in the original structure. Therefore, in contrast to that of a typical molluscan shell, the repair process of the eggcase was apparently identical to eggcase formation, as confirmed by the presence of identical microarchitecture and similar continuous crystal growth patterns.

Calcified granule deposition during the early stages of exoskeleton repair is a noteworthy characteristic of exoskeleton repair in mollusks^55,56^, and was also observed in the repaired regions of the argonaut eggcase. These granules were abundant on and between the initially formed organic membranes in the repaired eggcase, resembling previously reported deposits (Figs. 5, 6). Vertical sections revealed radiating growth patterns originating from these calcified granules, which resemble the nucleation points in original eggcase formation. This suggests that the calcified granules in the repaired eggcase can serve as crystal nucleation points. Mount et al. (2004) reported that granules involved in calcification are associated with hemocytes. However, chemical compositions of the calcified particles in repaired region and the original region in the eggcases were identical, in contrast to previous reports of compositional differences, particularly regarding phosphorus^56^. These findings suggest that eggcase regeneration does not entirely follow distinct repair pathways but employs a mechanism similar to normal formation.

Our findings allow us to reexamine Villepreux-Power’s widely accepted conclusion that argonauts form their eggcases using the first dorsal arms, based on her repair experiments (Fig. 8A). In her study, she partially damaged eggcases and observed the animals reattaching fragments using their first dorsal arms. Her observation likely corresponds to what we describe as fragment-inserted repair (FIR; Fig. 8B). However, since not all perforating damage can be repaired through FIR, the argonauts apparently possess another mechanism, the hole-sealing repair (HSR), which plugs the perforated area with newly secreted eggcase-forming materials lacking the layered structure of original eggcases (Fig. 8C). Overall, our observations raise doubts about the direct involvement of the first dorsal arms in eggcase formation, as suggested by Villepreux-Power, whose conclusions were based solely on behavioral observation and are thus inconclusive. Our findings implied that the first dorsal arms can secrete organic material, as indicated by the presence of the outermost organic membrane, and by the organic material enveloping fragment-inserted repair (Fig. 7). Microstructural characteristics support this skepticism, as layers mature within a few tens of millimeters from the aperture, where growth primarily occurs (Fig. 7). This growth mode contrasts with the fact that the expansive, web-like first dorsal arms of argonauts can cover the entire outer surface of the eggcase. Based on our observations, we propose that the first dorsal arms primarily serve a supportive role in eggcase formation. Future studies involving detailed histological observations, experimental re-examinations, and spatial gene expression analyses are essential to test this hypothesis.

### A sequential model of eggcase formation and biomineralization

The formation of mineralized skeletal structures is a sequential process, with each phase producing distinct morphological features. We identified four stages of prismatic structure formation in the argonaut eggcase (Fig. 7, S8). Stage one begins with crystallization during the formation of organic microfibers, referred to as the nucleation phase. We also observed an organic-inorganic particle within the organic fibers, likely serving as the nucleation site (Fig. S8, S9). Nucleation occurs both on the organic surface and within the organic microfibers (Figs. 5E, S8). In stage two, we observed a radial growth originating from the crystal nucleation site, followed by growth perpendicular to the eggcase surface, which we designate as the stage three. Similar growth stages have been described in stony coral skeletons^58^ and the avian eggshells^59^. Both crystal formation stages progress through competitive growth, but crystal morphologies differ between stages two and three. In the second stage, the outlines of the crystals and organic matter are clearly defined, while in the third stage, they are less distinct. This transition in crystal shape suggests the influence of different secreted matrix proteins involved in each growth stage. In the final stage, an organic membrane forms over the crystal layers.

Our proposed four-stage formation process which includes nucleation within organic fibers, radial crystal growth, perpendicular crystal growth, and the final organic membrane covering, successfully describes the microstructure observed in argonaut eggcase. This resembles rapidly growing calcareous structures like fibrous calcite in avian eggshells and fibrous aragonite in stony corals^60,61,62,63^. This resemblance suggests a possible convergence in microstructural strategies optimized for rapid growth. Stony corals with fibrous aragonite skeletal structures, such as *Acropora*, are among the fastest-growing corals, with a growth rate of up to 10 cm year^-1^ ^60^. Similarly, avian eggshells form in the oviduct at an approximate growth rate of 15 cm year^-1^ (0.02 mm h^-1^)^63^. The vertical growth rate of the microstructure during argonaut eggcase formation remains unknown, but it has been reported that argonaut eggcases undergo rapid growth towards the adoral area, reaching a maximum length of 17.4 cm in 4-5 months^64^. Such convergence may reflect the functional demands of efficiently forming robust calcareous structures under biological and environmental constraints. Understanding these processes may shed light on the evolution of extended phenotypes like the argonaut eggcase.

### The argonaut eggcase as an extended phenotype

Being a structure independent from the argonaut’s body but functionally integrated into the organism’s reproductive and ecological strategies^10,22^, the argonaut eggcase may be considered an extended phenotype^65^. Although its formation is behaviorally controlled^1,8^, the eggcase’s features like morphology, timing of production, and repair mechanisms, are consistent within species^14^, suggesting a degree of heritable regulation^13,15,23^. Its design is analogous to structures shaped by evolutionary selection acting on internal physiology or organism-mediated environmental modifications^66,67,68,69^. Like other examples of extended phenotypes such as beaver dams^67^ or spider webs^70,71^, the eggcase contributes directly to the organism’s fitness by facilitating buoyancy, pelagic reproduction, and egg protection^1,27^. Although not homologous to the molluscan shell^13,21^, the eggcase plays multiple organism-level roles and therefore could be interpreted as a behaviorally constructed external phenotype shaped by natural selection for the benefit of the replicator^65,66,71,72^. This may represent an example of the extended evolutionary synthesis (EES^73^), where adaptation to pelagicity might have induced the evolutionary trajectory to reinvent a shell-like mineralized structure as an extended phenotype^15^. In these contexts, the argonaut eggcase illustrates how behavioral and ecological contexts can convergently drive the evolution of functional and structural complexity.

## Materials and Methods

### Sample Collections

We obtained the eggcases of two argonaut species, *Argonauta argo* and *Argonauta hians*, through sampling collection and gifts (Table S1). The main samples for microstructural observation were collected from fixed nets in Kochi and Shimane prefectures in Japan, by local fishermen. Additional samples were donated by private contributors (Table S1). To ensure microstructural observation was conducted in a fresh state, samples were stored at −20 °C and analyzed within three months of collection. Technical terms were described in Supplementary Methods and Table S2.

### Scanning electron microscope (SEM) observation of argonaut eggcase microstructure

The microstructural observations were performed using a scanning electron microscope (SEM) VE-8800 (Keyence, Osaka, Japan). For preparation, samples were fragmented into smaller pieces using a precision cut-off machine TS-45 (Maruto Testing Machine Co., Tokyo, Japan). The observation surface was polished using the FG-18 polisher (Ryobi Ltd., Hiroshima, Japan), and then treated with 1M NaOH for 10 minutes to remove residual organic matter from the observation surface. The polished section was etched with Mutvei’s solution for 5 min^74^. The Mutvei’s solution, which consists of 500 ml 1% acetic acid, 500 ml 25% glutaraldehyde, and 10 g alcian blue powder, was used to etch biogenic carbonates and calcium phosphates, fix soluble and insoluble organic matrices, and stain mucopolysaccharides. After ultrasonic cleaning (US Cleaner USD-2R; AS ONE Corp., Osaka, Japan) for 10 minutes, the samples were air-dried overnight. Some samples were also observed under the stereoscopic microscope. The observation surface was coated with osmium using an osmium coater HPC-1SW (Vacuum Device Inc., Ibaraki, Japan). SEM imaging was performed with an accelerating voltage of 10-15 kV. The SEM stubs were vouchered at The University Museum, The University of Tokyo (Table S1).

### Scanning electron microscope and energy dispersive X-ray microanalysis (SEM-EDS)

Vertical sections of the eggcase and interior surfaces of the repair regions were prepared to identify the components and boundaries between organic and mineral regions, and to detect the elemental composition of the minerals. All samples were observed under SEM, with elemental compositions being analyzed by EDS at 15 kV using JSM-6010PLUS/LA (JEOL Ltd., Tokyo, Japan).

### Raman spectrometry

The crystal polymorph of calcium carbonate in the argonaut eggcase was identified using Raman spectroscopy. The eggcase was ground using a mortar and pestle before being placed on the sample stand. The samples were excited by an Ar ion laser (514.5 nm, 5500A, Ion Laser Technology, Salt Lake City, UT, USA). Raman spectra were obtained using a single polychromator (Chromex, 500is) fitted with a 1200 lines/mm grating and coupled to a charge-coupled device (CCD) camera with a resolution of 1024 × 128 pixels (Andor Technology, DU401A BR-DD).

## Data availability

All samples in this study were stored in The University Museum, The University of Tokyo. All data and electronic supplementary material are publicly available (uploded Supplementary Information).

## Acknowledgments

All authors thank Mitsu Ooshiki Co., Ltd. for their assistance in sample acquisition. We also thank Mr. Minoru Yoshida and his colleagues at Yoshida Suisan in the Oki Islands for their contribution to providing the sample.

DHES and KH thank past and present members of the Setiamarga Lab at NITW and Sasaki Lab at The University of Tokyo for their supports during the commencement of this study.

This study was partially supported by Takeda Science Foundation Life Science Research Grants 2022, Grants-in-aid for Basic Research KIBAN-C No. 19K12424 and 23K11511, both awarded to DHES, and Grants-in-aid for Basic Research KIBAN-C) No. 22K06340 awarded to MAY and DHES. KH was supported by JST SPRING, Grant No. JPMJSP2108 and the Sasakawa Scientific Research Grant from The Japan Science Society. KH was also supported by KOSEN GEAR 5.0 of National Institute of Technology: Agriculture and Fisheries Project awarded to DHES.

## Author contributions

KH, TS, and DHES conceived the study. TS and DHES supervised the project. KH conducted the main microstructural analyses, assisted by TS, TY, SO, HH, and TT. KH and MAY collected samples and biological data collection. KH wrote the first draft of the manuscript and KH, TS, and DHES edited subsequent versions of the manuscript. DHES wrote the final version of the manuscript. All authors were involved in data interpretation and discussions. All authors read, edited, and confirmed the content of the final version of the manuscript.

## Generative AI tools usage for language editing during manuscript preparation

During manuscript preparation, AI tools (mainly ChatGPT) were used to assist minor linguistic edits, such as grammar and stylistic adjustments, but not for content creation. Before using these tools, the manuscript was entirely prepared and written by the authors, mainly the first, second, and last authors, and further edited by all co-authors. After using these tools, the authors reviewed and edited the content as needed and take full responsibility for the content of the publication.

## Supplementary Information

### Supplementary Materials

*Terminology of orientation of the Argonaut Eggcase*

*Terminology of eggcase microstrcuture*

### Supplementary Tables

**Table S1 Samples for microstructural observation**

**Table S2 Technical terms of the eggcase microstructure of argonauts**

### Supplementary Figures

**Fig. S1 Raman spectrometry of calcified prismatic layer**

Three peaks indicate calcite of CaCO3 crystal polymorph.

**Fig. S2 EDS compositional analysis of the argonaut eggcase**

A) The chemical composition of the *A. argo* eggcase is shown in the spectrum. B) EDS elemental maps of the general microstructure of the eggcase of *A. argo*, showing a vertical section (Fig. 2B). A) SEM micrograph of the general microstructure of the eggcase of *A. argo*. C-G) Elemental maps of C) calcium, D) carbon, E) oxygen, F) sulfur, and G) agnesium.

Abbreviations: outer spherulitic-fibrous prismatic layer (o-sfpl), middle organic layer (m-ol), inner spherulitic-fibrous prismatic layer (i-sfpl)

**Fig. S3 Schematic diagrams, Stereomicroscopic images, and SEM micrographs of vertical sections of the *A. hians* eggcase**

A) General microstructure of the *A. hians* eggcase. A2, A3) Enlarged vertical secitons from A1. White arrowheads indicate boundary between crystals. B, C) Vertical section on two local regions: B) the center of the spiral and C) the keel tubercle. B2, B3) Enlarged vertical section from B1. C2) Enlarged vertical section of C1. C3) Vertical section of an obtuse keel tubercle. In the case of an obtuse angle, the middle organic layer is only curved without forming a significant gap.

Abbreviations: outer spherulitic-fibrous prismatic layer (o-sfpl), middle organic layer (m-ol), inner spherulitic-fibrous prismatic layer (i-sfpl)

**Fig. S4 Schematic diagrams and SEM micrographs of surface view of the *A. hians* eggcase** The microstructure of A-C) the interior surface and D-F) the exterior surface of the eggcase were captured at three regions: A, D) marginal, B, E) middle, and C, F) central regions. White arrowheads in A1 indicate the eggcase aperture. All samples for surface observation were unetched.

Abbreviations: crystallization termination floor (CTF)

**Fig. S5 Stereomicroscopic images of surface view of the *A. argo* and the *A. hians* eggcase** White arrowheads indicate the eggcase aperture. Green arrowheads indicate a tuberculate micro-ornamentation. Yellow arrowheads indicate the organic membrane.

**Fig. S6 SEM micrographs of vertical section of tuberculate micro-ornamentation of the *A. argo* eggcase**

**Fig. S7 SEM micrographs of EDTA eachted eggcase microstructure of *A. argo***

The organic fibers showed a reticulated growth pattern resembling that observed in the middle organic layer of the non-repaired eggcase following EDTA etching. A) SEM image showing the interior surface views of the repaired eggcase. B, C) SEM micrographs of the middle organic layer of the non-repaired eggcase after EDTA etching.

**Fig. S8 Schematic diagrams of the formation process of the spherulitic-fibrous prismatic structure in argonaut eggcase**

**Fig. S9 SEM Micrographs of the the nucleation site within the calcified layer of the *A. argo* eggcase**

A) SEM micrograph of a vertical section at the center of the spiral of the eggcase. B) Enlarged view of the edge of A. These micrographs indicate the occurrence of the nucleation site within the calcified layer, not limited to on the middle organic layer. White arrows indicate the nucleation site.

**Supplementary References**

## References

1. Finn, J. K. (2018). Recognising variability in the shells of argonauts (Cephalopoda: Argonautidae): the key to resolving the taxonomy of the family. Memoirs of Museum Victoria, 77, 63–104. 10.24199/j.mmv.2018.77.05

2. Taite, M., Fernández-Álvarez, F. Á., Braid, H. E., Bush, S. L., Bolstad, K., Drewery, J., … & Allcock, A. L. (2023). Genome skimming elucidates the evolutionary history of Octopoda. Molecular Phylogenetics and Evolution, 182, 107729. 10.1016/j.ympev.2023.107729

3. Bello, G. (2012). Exaptations in argonautoidea (cephalopoda: Coleoidea: Octopoda). Neues Jahrbuch fur Geologie und Palaontologie–Abhandlungen, 266(1), 85. 10.1127/0077-7749/2012/0290

4. Setiamarga, D. H., Hirota, K., Yoshida, M. A., Takeda, Y., Kito, K., Ishikawa, M., … & Endo, K. (2021). Hydrophilic shell matrix proteins of *Nautilus pompilius* and the identification of a core set of conchiferan domains. Genes, 12(12), 1925. 10.3390/genes12121925

5. Hirota, K., Tochino, N., Seto, M., Sasaki, T., Yoshida, M. A., & Setiamarga, D. H. (2023). Comparative proteomics of the shell matrix proteins of *Nautilus pompilius* and the conchiferans provide insights into mollusk shell evolution at the molecular level. Marine Biology, 170(9), 106. 10.1007/s00227-023-04244-x

6. Finn, J. K., & Norman, M. D. (2010). The argonaut shell: gas–mediated buoyancy control in a pelagic octopus. Proceedings of the royal society b: Biological sciences, 277(1696), 2967–2971. 10.1098/rspb.2010.0155

7. Naef, A. (1923) Cephalopoda (systematics). Fauna and Flora of the Bay of Naples, Monograph 35, part 1, vol. 1, pp. 293–917. Translated from German. Jerusalem, Israel: Israel Program for Scientific Translations.

8. Villepreux-Power, J. (1856). Observations physiques sur le poulpe de l’Argonauta argo: commencées en 1832 et terminées en 1843, dédiées à M. le professeur Owen FRS. Impr. Charles de Mourges frères.

9. Strugnell, J., Jackson, J., Drummond, A. J., & Cooper, A. (2006). Divergence time estimates for major cephalopod groups: evidence from multiple genes. Cladistics, 22(1), 89–96. 10.1111/j.1096-0031.2006.00086.x

10. Strugnell, J., & Allcock, A. L. (2010). Co–estimation of phylogeny and divergence times of Argonautoidea using relaxed phylogenetics. Molecular Phylogenetics and Evolution, 54(3), 701–708. 10.1016/j.ympev.2009.11.017

11. Sanchez, G., Setiamarga, D. H., Tuanapaya, S., Tongtherm, K., Winkelmann, I. E., Schmidbaur, H., … & Nabhitabhata, J. (2018). Genus-level phylogeny of cephalopods using molecular markers: current status and problematic areas. PeerJ, 6, e4331. 10.7717/peerj.4331

12. Hirota, K., Yoshida, M. A., Itoh, T., Toyoda, A., & Setiamarga, D. H. (2021). The full mitochondrial genome sequence of the greater argonaut *Argonauta argo* (Cephalopoda, Argonautoidea) and its phylogenetic position in Octopodiformes. Mitochondrial DNA Part B, 6(4), 1451–1453. 10.1080/23802359.2021.1911710

13. Oudot, M., Shir, I. B., Schmidt, A., Plasseraud, L., Broussard, C., Neige, P., & Marin, F. (2020). A nature’s curiosity: the argonaut “shell” and its organic content. Crystals, 10(9), 839. 10.3390/cryst10090839

14. Checa, A. G., Linares, F., Grenier, C., Griesshaber, E., Rodríguez-Navarro, A. B., & Schmahl, W. W. (2021). The argonaut constructs its shell via physical self–organization and coordinated cell sensorial activity. Iscience, 24(11). 10.1016/j.isci.2021.103288

15. Yoshida, M. A., Hirota, K., Imoto, J., Okuno, M., Tanaka, H., Kajitani, R., … & Setiamarga, D. H. (2022). Gene recruitments and dismissals in the argonaut genome provide insights into pelagic lifestyle adaptation and shell–like eggcase reacquisition. Genome Biology and Evolution, 14(11), evac140. 10.1093/gbe/evac140

16. Cuif, J. P., Lecointre, G., Perrin, C., Tillier, A., & Tillier, S. (2003). Patterns of septal biomineralization in Scleractinia compared with their 28S rRNA phylogeny: a dual approach for a new taxonomic framework. Zoologica Scripta, 32(5), 459–473. 10.1046/j.1463-6409.2003.00133.x

17. Sato, K., Kano, Y., Setiamarga, D. H., Watanabe, H. K., & Sasaki, T. (2020). Molecular phylogeny of protobranch bivalves and systematic implications of their shell microstructure. Zoologica Scripta, 49(4), 458–472. 10.1111/zsc.12419.

18. Meldrum, F. C. (2003). Calcium carbonate in biomineralisation and biomimetic chemistry. International Materials Reviews, 48(3), 187–224. 10.1179/095066003225005836

19. Rodriguez-Navarro, A., Jimenez-Lopez, C., Hernandez-Hernandez, A., Checa, A., & García-Ruiz, J. M. (2008). Nanocrystalline structures in calcium carbonate biominerals. Journal of Nanophotonics, 2(1), 021935. 10.1117/1.3062826

20. Kobayashi, I. (1971). Internal microstructure of the shell of *Argonauta argo*. Venus, 30, 103–112. 10.18941/venusjjm.30.3_103

21. Mitchell, P. R., Phakey, P. P., and Rachinger, W. A. (1994). Ultrastructural observations of the argonaut shell. Scanning microscopy 8(1), 4. 10.18941/venusjjm.30.3_103

22. Wolfe, K., Smith, A. M., Trimby, P., & Byrne, M. (2013). Microstructure of the paper nautilus (*Argonauta nodosa*) shell and the novel application of electron backscatter diffraction (EBSD) to address effects of ocean acidification. Marine biology, 160, 2271–2278. 10.1007/s00227-012-2032-4.

23. Kniprath, E. (1981). Ontogeny of the molluscan shell field: a review. Zoologica Scripta, 10(1), 61–79. 10.1111/j.1463-6409.1981.tb00485.x

24. Sukhsangchan, C., & Nabhitabhat, J. (2007). Embryonic development of muddy paper nautilus, *Argonauta hians* Lightfoot, 1786, from Andaman Sea, Thailand. Agriculture and Natural Resources, 41(3), 531–538.

25. Wolfe, K., Smith, A. M., Trimby, P., & Byrne, M. (2012). Vulnerability of the paper nautilus (*Argonauta nodosa*) shell to a climate–change ocean: potential for extinction by dissolution. The Biological Bulletin, 223(2), 236–244. 10.1086/BBLv223n2p236

26. Finn, J. K. (2013). Taxonomy and biology of the argonauts (Cephalopoda: Argonautidae) with particular reference to Australian material. Molluscan Research, 33(3), 143–222. 10.1080/13235818.2013.824854

27. Lemanis, R., Tadayon, K., Reich, E., Joshi, G., Johannes Best, R., Stevens, K., & Zlotnikov, I. (2022). Wet shells and dry tales: the evolutionary ‘Just–So’stories behind the structure–function of biominerals. Journal of the Royal Society Interface, 19(191), 20220336. 10.1098/rsif.2022.0336

28. Jaya, B. N., Hoffmann, R., Kirchlechner, C., Dehm, G., Scheu, C., & Langer, G. (2016). Coccospheres confer mechanical protection: New evidence for an old hypothesis. Acta Biomaterialia, 42, 258–264. 10.1016/j.actbio.2016.07.036

29. D’Alba, L., Torres, R., Waterhouse, G. I., Eliason, C., Hauber, M. E., & Shawkey, M. D. (2017). What does the eggshell cuticle do? A functional comparison of avian eggshell cuticles. Physiological and Biochemical Zoology, 90(5), 588–599. 10.1086/693434

30. Walker, C. E., Heath, S., Salmon, D. L., Smirnoff, N., Langer, G., Taylor, A. R., … & Wheeler, G. L. (2018). An extracellular polysaccharide–rich organic layer contributes to organization of the coccosphere in coccolithophores. Frontiers in Marine Science, 5, 306. 10.3389/fmars.2018.00306

31. Walker, J. M., & Langer, G. (2021). Coccolith crystals: Pure calcite or organic–mineral composite structures?. Acta Biomaterialia, 125, 83–89. 10.1016/j.actbio.2021.02.025

32. Kulshreshtha, G., D’alba, L., Dunn, I. C., Rehault–Godbert, S., Rodriguez–Navarro, A. B., & Hincke, M. T. (2022). Properties, genetics and innate immune function of the cuticle in egg–laying species. Frontiers in Immunology, 13, 838525. 10.3389/fimmu.2022.838525

33. Marin, F., Le Roy, N., & Marie, B. (2012). The formation and mineralization of mollusk shell. Frontiers in Bioscience, 4(3), 1099–1125. 10.2741/s321

34. Sprey, A. M. (2001). Tuberculate micro-ornament on the juvenile shell of Middle Jurassic ammonoids. Lethaia, 34(1), 31–35. 10.1080/002411601300068215

35. Dauphin, Y., Luquet, G., Percot, A., & Bonnaud–Ponticelli, L. (2020). Comparison of embryonic and adult shells of *Sepia officinalis* (Cephalopoda, Mollusca). Zoomorphology, 139, 151–169. 10.1007/s00435-020-00477-2

36. Checa, A. G., Grenier, C., Griesshaber, E., Schmahl, W. W., Cartwright, J. H., Salas, C., & Oudot, M. (2022). The shell structure and chamber production cycle of the cephalopod Spirula (Coleoidea, Decabrachia). Marine Biology, 169(10), 132. 10.1007/s00227-022-04120-0

37. Bandel, K. (1985). Cephalopod morphology and function. Series in Geology, Notes for Short Course, 13, 190–201. 10.1017/S0271164800001172

38. Saunders, W. B., & Landman, N. (Eds.). (2009). Nautilus: the biology and paleobiology of a living fossil, reprint with additions (Vol. 6). Springer Science & Business Media.

39. Landman, N. H., Tanabe, K., & Davis, R. A. (Eds.). (2013). Ammonoid paleobiology (Vol. 13). Springer Science & Business Media.

40. Huang, J., & Zhang, R. (2022). The mineralization of molluscan shells: Some unsolved problems and special considerations. Frontiers in Marine Science, 9, 874534. 10.3389/fmars.2022.874534

41. Kobayashi, T. (1954). A new Palaeogene paracenoceratoid from southern Kyushu in Japan. Japanese Journal of Geology and Geography, 24, 181–184.

42. Ries, J. B. (2009). Effects of secular variation in seawater Mg/Ca ratio (calcite–aragonite seas) on CaCO3 sediment production by the calcareous algae Halimeda, Penicillus and Udotea–evidence from recent experiments and the geological record. Terra Nova, 21(5), 323–339. 10.1111/j.1365-3121.2009.00899.x

43. Doguzhaeva, L. A., Weaver, P. G., & Ciampaglio, C. N. (2012). A unique late Eocene coleoid cephalopod Mississaepia from Mississippi, USA: New data on cuttlebone structure, and their phylogenetic implications. Acta Palaeontologica Polonica, 59(1), 147–162. 10.4202/app.2011.0208

44. Arkhipkin, A. I., Bizikov, V. A., & Fuchs, D. (2012). Vestigial phragmocone in the gladius points to a deepwater origin of squid (Mollusca: Cephalopoda). Deep Sea Research Part I: Oceanographic Research Papers, 61, 109–122. 10.1016/j.dsr.2011.11.010

45. Tanabe, K., Kulicki, C., & Landman, N. H. (2008). Development of the embryonic shell structure of Mesozoic ammonoids. American Museum Novitates, 2008(3621), 1–19. 10.1206/588.1

46. Bandel, K., Landman, N. H., & Waage, K. M. (1982). Micro–ornament on early whorls of Mesozoic ammonites: implications for early ontogeny. Journal of Paleontology, 5(1), 386–391.

47. Landman, N. H., Bizzarini, F., Tanabe, K., Mapes, R. H., & Kulicki, C. (2001). Micro– ornamentation on the embryonic and postembryonic shell of *Triassic ceratites* (Ammonoidea). American Malacological Bulletin, 16(1/2), 1–12.

48. Hickman, C. S. (2004). The problem of similarity: analysis of repeated patterns of microsculpture on gastropod larval shells. Invertebrate Biology, 123(3), 198–211. 10.1111/j.1744-7410.2004.tb00155.x

49. Tanabe, K, Kulicki, C, Landman, N. H, & Kaim, A. (2010). Tuberculate micro– ornamentation on embryonic shells of Mesozoic ammonoids: microstructure, taxonomic variation, and morphogenesis. Cephalopods–Present and Past, 105–125.

50. Landman, N. H. (1994). Exceptionally well–preserved ammonites from the Upper Cretaceous (Turonian–Santonian) of North America: implications for ammonite early ontogeny. American Museum novitates; no. 3086.

51. Saleuddin, A. S. M., & Wilbur, K. M. (1969). Shell regeneration in *Helix pomatia*. Canadian journal of zoology, 47(1), 51–53. 10.1139/z69-011

52. Meenakshi, V. R., Blackwelder, P. L., & Wilbur, K. M. (1973). An ultrastructural study of shell regeneration in *Mytilus edulis* (Mollusca: Bivalvia). Journal of Zoology, 171(4), 475–484. 10.1111/j.1469-7998.1973.tb02229.x

53. Meenakshi, V. R., Martin, A. W., & Wilbur, K. M. (1974). Shell repair in *Nautilus macromphalus*. Marine Biology, 27, 27–35. 10.1007/BF00394757

54. Su, X. W., Zhang, D. M., & Heuer, A. H. (2004). Tissue regeneration in the shell of the giant queen conch, *Strombus gigas*. Chemistry of Materials, 16(4), 581–593. 10.1021/cm030573l

55. Cho, S. M., Lee, Y. M., & Jeong, W. G. (2011). Crassostrea gigas. Korean J. Malacol, 27(1), 35–42. 10.9710/kjm.2011.27.3.223

56. Li, S., Liu, Y., Liu, C., Huang, J., Zheng, G., Xie, L., & Zhang, R. (2016). Hemocytes participate in calcium carbonate crystal formation, transportation and shell regeneration in the pearl oyster *Pinctada fucata*. Fish & shellfish immunology, 51, 263–270. 10.1016/j.fsi.2016.02.027

57. Mount, A. S., Wheeler, A. P., Paradkar, R. P., & Snider, D. (2004). Hemocyte-mediated shell mineralization in the eastern oyster. Science, 304(5668), 297–300. 10.1126/science.1090506

58. Cuif, J. P., Dauphin, Y., & Gautret, P. (1997). Biomineralization features in scleractinian coral skeletons: source of new taxonomic criteria. Boletín de la Real Sociedad Española de Historia Natural (Sección Geológica), 92, 129–141.

59. Gautron, J., Stapane, L., Le Roy, N., Nys, Y., Rodriguez–Navarro, A. B., & Hincke, M. T. (2021). Avian eggshell biomineralization: an update on its structure, mineralogy and protein tool kit. BMC Molecular and Cell Biology, 22, 1–17. 10.1186/s12860-021-00351-z

60. Lirman, D. (2000). Fragmentation in the branching coral Acropora palmata (Lamarck): growth, survivorship, and reproduction of colonies and fragments. Journal of Experimental Marine Biology and Ecology, 251(1), 41–57. 10.1016/S0022-0981(00)00205-7

61. Nothdurft, L. D., & Webb, G. E. (2007). Microstructure of common reef–building coral genera Acropora, Pocillopora, Goniastrea and Porites: constraints on spatial resolution in geochemical sampling. Facies, 53, 1–26. 10.1007/s10347-006-0090-0

62. Takeuchi, T., Yamada, L., Shinzato, C., Sawada, H., & Satoh, N. (2016). Stepwise evolution of coral biomineralization revealed with genome–wide proteomics and transcriptomics. PLoS One, 11(6), e0156424. 10.1371/journal.pone.0156424

63. Karlsson, O., & Lilja, C. (2008). Eggshell structure, mode of development and growth rate in birds. Zoology, 111(6), 494–502. 10.1016/j.zool.2007.11.005

64. Stevens, K., Iba, Y., Suzuki, A., & Mutterlose, J. (2015). Biological and environmental signals recorded in shells of *Argonauta argo* (Cephalopoda, Octobrachia) from the Sea of Japan. Marine Biology, 162, 2203–2215. 10.1007/s00227-015-2750-5

65. Dawkins, R. (1982). The extended phenotype (Vol. 8). Oxford: Oxford university press.

66. Turner, J. S. (2009). The extended organism: the physiology of animal-built structures. Harvard University Press.

67. Dawkins, R. (2004). Extended phenotype–but not too extended. A reply to Laland, Turner and Jablonka. Biology and Philosophy, 19, 377–396. 10.1023/b:biph.0000036180.14904.96

68. Japyassú, H. F., & Laland, K. N. (2017). Extended spider cognition. Animal Cognition, 20(3), 375–395. https://doi-org.utokyo.idm.oclc.org/10.1007/s10071-017-1069-7

69. Hunter, P. (2018). The revival of the extended phenotype: After more than 30 years, Dawkins’ extended phenotype hypothesis is enriching evolutionary biology and inspiring potential applications. EMBO reports, 19(7), e46477. 10.15252/embr.201846477

70. Nakata, K. (2012). Plasticity in an extended phenotype and reversed up-down asymmetry of spider orb webs. Animal Behaviour, 83(3), 821–826. 10.1016/j.anbehav.2011.12.030

71. Blamires, S. J., Martens, P. J., & Kasumovic, M. M. (2018). Fitness consequences of plasticity in an extended phenotype. Journal of Experimental Biology, 221(4), jeb167288. 10.1242/jeb.167288

72. Hunter, P. (2009). Extended phenotype redux: How far can the reach of genes extend in manipulating the environment of an organism?. EMBO reports, 10(3), 212–215. 10.1038/embor.2009.18

73. Laland, K. N., Uller, T., Feldman, M. W., Sterelny, K., Müller, G. B., Moczek, A., … & Odling-Smee, J. (2015). The extended evolutionary synthesis: its structure, assumptions and predictions. Proceedings of the royal society B: biological sciences, 282(1813), 20151019. 10.1098/rspb.2015.1019

74. Schöne, B. R., Dunca, E., Fiebig, J., & Pfeiffer, M. (2005). Mutvei’s solution: an ideal agent for resolving microgrowth structures of biogenic carbonates. Palaeogeography, Palaeoclimatology, Palaeoecology, 228(1-2), 149–166. 10.1016/j.palaeo.2005.03.054

